# DISRUPTION OF SYMPATHETIC OUTFLOW TO INTRA-ABDOMINAL ORGANS EMULATES THE METABOLIC EFFECTS OF SLEEVE GASTRECTOMY IN OBESE MICE

**DOI:** 10.1101/2022.10.17.512615

**Authors:** Gary J. Schwartz, Rogerio O. Batista, Natalie R. Lopatinsky, Marko Kraljević, Caroline S. Jiang, Amanda S. Dirnberger, Ana B. Emiliano

## Abstract

Although sleeve gastrectomy (SG) is the most commonly performed bariatric surgery in the US, its mechanistic underpinnings have not been fully determined. Thus, we set out to investigate whether SG’s effects on the peripheral sympathetic system could mediate the metabolic effects of SG. The celiac-superior mesenteric ganglia (CSMG) lie juxtaposed to the stomach and supply the sympathetic innervation of the stomach, as well as to numerous intra-abdominal organs relevant to metabolism. Here we investigated the effects of SG on the CSMG. SG led to the degeneration of neurons in the CSMG, as evidenced by chromatolysis, which was not found in control mice. Furthermore, CSMG ablation (CGX) completely recapitulated the glycemic and weight loss effects of SG, promoting weight loss at the expense of fat mass in both males and females. Glycemic improvement was robust in males but much more modest in female mice. Norepinephrine tissue content measurement by high performance liquid chromatography revealed that liver, duodenum, and ileum were organs where both SG and CGX displayed evidence of significant sympathetic denervation. Both SG and CGX were associated with increased levels of glucagon-like peptide 1 (GLP-1) and high free fatty acid content in the stools. In conclusion, CSMG neuronal degeneration caused by SG appears to be a mediator of the metabolic effects of this type of bariatric surgery.

## INTRODUCTION

Despite the popularity of bariatric surgery as a treatment for obesity and type 2 diabetes mellitus (T2DM), the exact mechanisms through which bariatric surgery leads to glycemic improvement and durable weight loss are largely unknown. The potential involvement of the vagal motor and sensory innervation of the stomach and other parts of the gastrointestinal (GI) tract in the achievement of the beneficial metabolic outcomes of bariatric surgery has garnered substantial attention in the preclinical literature, especially in relation to RYGB (1–5). The rationale for this emphasis on studying the effects of bariatric surgery on the vagal innervation stem in part from the rich vagal sensory motor innervation of the stomach (6) and the fact that vagal sensory terminals have the important function of detecting changes in pressure, volume and nutrient exposure (7). However, the stomach also receives rich sympathetic innervation, mainly from the celiac-superior mesenteric ganglia (CSMG) (8). The CSMG are prevertebral sympathetic ganglia classically believed to transduce central nervous system (CNS) input to the target organs it innervates, which include liver, pancreas, stomach, intestines, kidneys, and adrenal glands (9–12). Much less is known about the potential role played by sympathetic CSMG neurons in obesity or metabolic disease. Sympathetic nervous system (SNS) overactivity is a hallmark of obesity accompanied by the metabolic syndrome (13). Weight loss, either through lifestyle modifications or sleeve gastrectomy (SG), has been shown to reduce SNS activity, while also decreasing cardiovascular mortality and improving glucose homeostasis (14, 15). Increased SNS activity has been linked to insulin resistance (16, 17) and negative clinical outcomes in heart failure, kidney disease, coronary artery disease and stroke (18–21). A recent study pointed to the potential involvement of CSMG neurons in glucose homeostasis via input received from cocaine- and amphetamine-regulated transcript^+^(CART) viscerofugal neurons in the intestine (22). CSMG neurons have also been implicated in the sympathoadrenal response to hypoglycemia, as CSMG ganglionectomy (CGX) in rats led to a 50% decrease in the epinephrine and norepinephrine response to insulin-induced hypoglycemia (23). It is not known, however, whether sympathetic ganglia are mediators of the deleterious effects imparted by sympathetic overactivity on cardiovascular and metabolic function in obesity. Furthermore, although a decrease in leptin and insulin plasma levels that results from weight loss are thought to be linked to the SNS decline in activity after any type of weight loss, the exact mechanisms through which weight loss promotes SNS inhibition remain largely undetermined. Moreover, it is not known if SG decreases SNS activity through direct, weight loss independent mechanisms.

Consequently, using 18-week-old wild type, diet-induced obese (DIO) mice, we set out to investigate whether SG exerted any effects on the CSMG and whether these effects could be related to SG-induced blood glucose lowering and weight loss.

## Material and Methods

### Animals, Housing and Experimental Design

The studies were performed according to ARRIVE guidelines and in accordance with the Columbia University Institutional Animal Care and Use Committee, which granted approval for all the experiments in this study. Female C57bl/6J mice were purchased from The Jackson Laboratories at 6 weeks of age and maintained thereafter on a high fat diet (Research Diets, catalogue number D12492, New Jersey, NJ). C57bl/6J obese female mice are not commercially available. Male C57bl/6J raised on a 60% high fat diet from 6 weeks of age, were purchased from The Jackson Laboratories at 15-16 weeks of age and maintained on a high fat diet throughout all studies. Mice were group housed in a vivarium maintained at 22°C on a 12:12 (7 am - 7 pm) light / (7 pm - 7 am) dark cycle. At 18 weeks of age, mice were randomized to SG (sleeve gastrectomy), CGX (celiac ganglionectomy), SH-IF (sham surgery weight matched to the SG group through calorie restriction and intermittent fasting) and SH-AL (sham surgery with ad libitum feeding) (n=12-14 for SG; n=8-12 for SH-IF, n=8-12 for SH-AL, for both male and female cohorts). Intermittent fasting (IF) involved feeding the SH-IF group a variable ration, once daily, at different times of the day, on weekdays, with the amount depending on their weight loss or gain, so that they would be weight-matched to the SG group. From Fridays to Mondays, SH-IF mice were allowed to eat without restriction (*ad libitum*). Baseline measurements were defined as having been taken approximately 2 weeks prior to surgeries. Body weight (BW) and food intake were measured daily for the first week post-operatively and weekly thereafter. Mice at 4 weeks post-surgery were placed under isoflurane anesthesia, after an overnight fast (8-10 hours), and intracardiac perfusion was performed with 0.9% saline, followed by 4% paraformaldehyde, and liver was harvested for clearing (iDisco, see below). For CSMG microscopy, the CSMG from mice that underwent SG, SH-AL and SH-IF were dissected, placed in 4% paraformaldehyde overnight and transferred to 30% sucrose for 2 days before being frozen for sectioning in a cryostat.

*Surgical Procedures*. Sleeve gastrectomy was performed in anesthetized mice (isoflurane) to remove approximately 80% of the stomach, as previously described previously (24, 25). For the sham surgery, a laparotomy was followed by isolation of the stomach, severance of the gastrosplenic and gastrophrenic ligaments, followed by exertion of pressure with a smooth forceps along a vertical line between the esophageal sphincter and the pylorus, at the point where the stomach is resected in the VSG procedure. The stomach was placed back into the abdominal cavity and the laparotomy closed in layers. CGX was performed by carefully dissecting all the nerve tissue lying between and around the celiac artery and superior mesenteric artery, and which also encroaches on the abdominal aorta (figure 1A).

**Figure 1.**
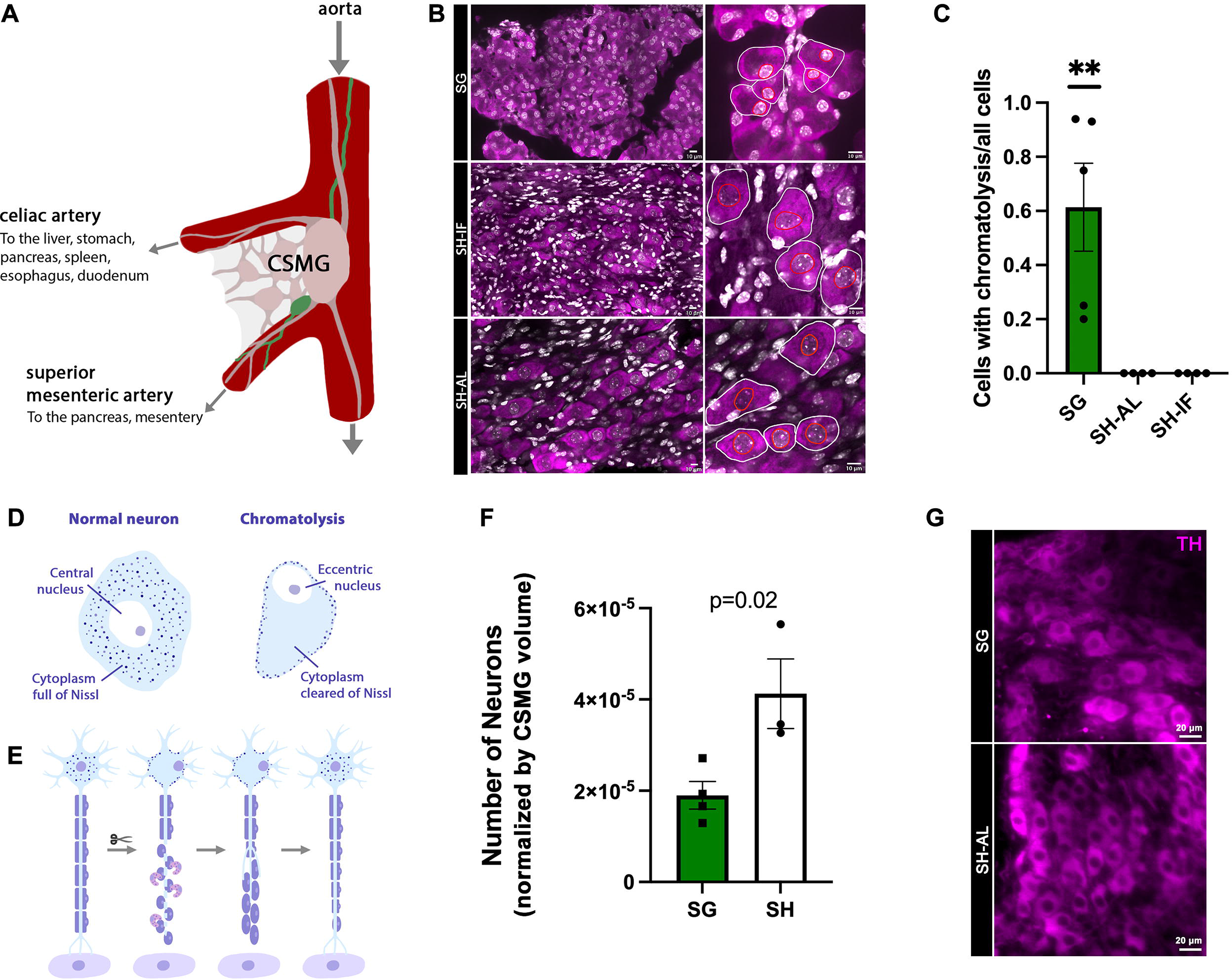
**A**. Anatomic boundaries of the celiac-superior mesenteric ganglia (CSMG) in the mouse. **B**. Confocal images of 20 micrometer-CSMG slices from mice that underwent SG (sleeve gastrectomy), sham surgery with caloric restriction and intermittent fasting (SH-IF), and sham surgery with ad libitum feeding (SH-AL). Chromatolysis was abundantly present in SG mice, while none was observed in SH-IF and SH-AL mice. Neurons were stained with NeuroTrace™ 640/660. Scale bar 10 μm. **C**. Percentage of neurons with chromatolysis over total number of neurons counted. Normalized by total number of neurons counted. One-way ANOVA. N=2-3 mice per group. **p<0.01. **D**. Schematic depicting chromatolysis. **E**. Schematic showing chromatolysis leading to denervation of a specific target organ, followed by recovery, which is a potential fate of neurons undergoing degenerative changes after axonal injury. **F**. Total neuronal cell count from 4 tissue-cleared CSMG from SG mice and 3 from SH-AL mice. Count was normalized by CSMG area and volume and performed with the use of the software Stereology™. Unpaired t-test. **G**. Light sheet images of tissue-cleared CSMG stained for tyrosine hydroxylase immunoreactivity, showing a significantly larger population of neurons in SH-AL mice compared to SG mice.

### Glucose Tolerance Test, Insulin Tolerance Test, Glucose Stimulated Insulin Secretion and Mixed Meal Test

Mice were fasted for 6 hours prior to glucose and insulin tolerance tests. SH-IF mice were fed from 0.8-1.0 g of high fat diet one hour prior to the start of the 6-hour fast, in preparation for the glucose and insulin tolerance tests. This was done to avoid a more prolonged period of fasting in SH-IF mice, compared to SG and SH-AL mice. Oral glucose tolerance tests were performed at baseline, and then at 1 and 3 weeks post-operatively; insulin tolerance tests were performed 2 and 4 weeks post-operatively. Oral glucose tolerance tests (OGTT) were performed by oral gavage using a plastic feeding tube (1FTP-20-38, Instech Laboratories, Inc.), with 2 grams of dextrose per kg of body weight. Insulin tolerance tests (ITT) were performed using regular insulin at a dose of 0.75 units per Kg of body weight, injected intraperitoneally (Humulin R, Lilly). Two weeks post-surgery, a cohort of mice underwent a glucose stimulated insulin secretion test an OGTT after an overnight fast. Insulin was measured at time points 0, 10, 20 and 30 minutes, from tail blood collection. A mixed meal test was performed by on a separate cohort of mice after an overnight fast, also 2 weeks post-surgery. An oral gavage with 200 μl of Ensure® was administered. Tail blood was collected at baseline and at 30 minutes after gavage.

### Urine Glucose Detection

Chemical strips (Jmd International™) with a lower limit of detection between 50-100 mg/dL of glucose were used to qualitatively detect the presence of glucose in the urine of male mice.

### Body Composition, Energy expenditure and Locomotor Activity

Body composition was measured at baseline, and weekly post-operatively using Echo-MRI™-100H (EchoMRI LLC, Houston, TX). Energy expenditure by indirect calorimetry was performed by singly housing mice in the Comprehensive Lab Animal Monitoring System (CLAMS, Columbus Instruments, Columbus, OH) for one week, between 3-4 weeks post-operatively. The SH-IF group was fed ad libitum during the calorimetry experiment. CLAMS measurements included Volume O_2_, O_2_ in and O_2_ out, delta O_2_, Accumulated O_2_, Volume CO_2_, CO_2_ in and CO_2_ out, delta CO_2_, Accumulated CO_2_, RER, Heat, Flow, Feed Weight, Feed Accumulation, and Total Locomotor Activity, which was calculated as the sum of activity along the X, Y and Z axes of the chambers.

### Fecal Free-Fatty Acid Content Assay

Feces were collected from individual mice at 1, 2 and 4 weeks post-operatively (they were not collected during the third week because the mice were undergoing indirect calorimetry). Feces were dried overnight on a six-well plate, at 42°C. For each reaction, 100 mg of feces (per mouse) were placed in 1 mL of NaCl and then homogenized with beads for five minutes. The solution was transferred to a 15 mL tube, to which a mix of chloroform and methanol at 2:1 was added (Chloroform, Fisher, C298-4; Methanol, Fisher A456-212). The mixture was vortexed vigorously and then centrifuged at 2000g for 10 minutes. The chloroform phase was collected and transferred to a 20 mL vial and evaporated under dinitrogen. After that, 1 mL of deionized water was added. The next day, samples were assayed in duplicates, with a Free Fatty Acid (FFA) kit (Wako 995-34693; 997-34893; Fujifilm USA). Samples were diluted at 1:20 and incubated at 37°C for five minutes and read with a microplate reader.

### Tissue Norepinephrine Content

Liver, pancreas, duodenum, jejunum, ileum, mesenteric white adipose tissue (mWAT), inguinal white adipose tissue (iWAT), gonadal white adipose tissue (gWAT) and interscapular brown adipose tissue (iBAT) were collected 4 weeks post-surgery from overnight fasted mice (8-10 hours) and immediately frozen on dry ice and transferred to a minus 80 freezer. Tissues were then homogenized for measurement of norepinephrine by high performance liquid chromatography, as previously described (26).

### Plasma Assays

At 2 weeks post-surgery, blood was collected from tail vein at baseline and after mice received an oral gavage with 200 μL of Ensure, and their blood was collected via cardiac puncture. Blood was placed in serum separator tubes (Microvette, Kent Scientific catalogue #MTSC-SER), to which were added aprotinin, diprotin A (a DPPIV inhibitor) and Halt protease inhibitor cocktail (aprotinin, Millipore Sigma, catalogue #A1153; diprotin A, Enzo Scientific, catalogue #ALX-260-036-M005; HALT, Thermo Scientific catalogue # PI87785). Blood was spun for 15 min at 2000 g in a refrigerated benchtop microcentrifuge and subsequently stored at −80°C. Serum insulin and active GLP-1 were measured using a U-PLEX MSD assay (MesoScale Discovery, catalogue # K152ACM-1).

### Cholera toxin B injections in liver, pancreas, and mesenteric white adipose tissue

Cholera toxin B (CTb) conjugated with Alexa fluor 647 and 555 were injected into the liver parenchyma (total of 10 μL split through liver segments) and pancreas (total of 10 μL divided into head, body, and tail), respectively (Invitrogen, 647, #C34778; Invitrogen, 555, # C34776). Then, in a separate experiment, CTb conjugated with Alexa Fluor 647 was injected into the mesenteric white adipose tissue (mWAT) (10 μL total). In a third experiment, mice were injected with Ctb conjugated with Alexa Fluor 488, 555 and 647 in pancreas, liver and mWAT, respectively, in the same manner as described above (CTb Invitrogen, 488, # C34775). Mice were euthanized 5 days after injections, their CSMG harvested, fixed overnight in fresh 4% paraformaldehyde, and transferred to 30% sucrose. Whole CSMG were processed for iDisco as previously described (27). Images were captured with a Ultramicroscope II Light sheet microscope (2×0.5NA objective, with a 1X zoom) and analyzed using Imaris software according (Imaris 9.2 Reference Manual).

### Immunohistochemistry and liver tissue clearing

For identification of chromatolysis, CSMG were dissected from mice 4 weeks post-surgery and kept in 4% paraformaldehyde (PFA) (Electron Microscopy Sciences) overnight. The next day, they were transferred to 30% sucrose in phosphate-buffered saline (PBS) where they remained for one week. Subsequently, they were flash frozen and sectioned in a cryostat at 20 μm sections. CSMG were sliced into 20 μm sections which were stained with the fluorescence Nissl stain NeuroTrace 640/660 Deep Red fluorescence (Invitrogen, catalogue #N21483) at a dilution of 1:150 and counterstained with DAPI (ThermoScientific, #62247). Prolong Diamond antifade mountant (Invitrogen, # P36970) was used for cover slipping. Images were taken with a spinning disk confocal microscope, at 20X magnification.

For the CSMG neuronal cell count, tissue clearing was performed and anti-TH antibody (Millipore, #AB152) was used to label sympathetic neurons in the CSMG. Images were captured with an Ultramicroscope II Light sheet microscope (2×0.5NA objective, with a 1X zoom) and analyzed using the software Stereology™ for precise neuronal cell counting with normalization for CSMG area and volume.

Liver iDisco was performed as previously described (28). Primary antibodies were anti-Protein Gene Product 9.5 (PGP9.5) (Invitrogen, #PA1-10011) and anti-Tyrosine Hydroxylase (TH) (Millipore, #AB152) at 1:200 dilution for both antibodies. Secondary antibodies were Alexa Fluor 647 Goat anti-Chicken (Invitrogen, #A-21449) and Alexa Fluor 790 Goat anti-Rabbit (Invitrogen, #A11369) at 1:200 dilution for both antibodies. Images were obtained using a Ultramicroscope II Light sheet microscope (2×0.5NA objective, with a 1X zoom) and analyzed using Imaris software according (Imaris 9.2 Reference Manual). iDisco images presented in this manuscript are raw data.

### Statistics

We used GraphPad Prism 9 version software. Non-parametric Mann Whitney tests were used for two-group comparisons of a single measure (student’s t test). One-way ANOVA with Kruskal-Wallis tests were used for comparisons of single measures among more than two groups. Two-way ANOVA with Multiple Comparisons with Tukey tests were used for analysis of multiple groups involving repeated measures. Column factor refers to surgical group and row factor to time. Data are expressed as means ± SEM. Area Under the Curve (AUC) was generated for OGTT and ITT and then compared using One-way ANOVA with Kruskal-Wallis tests. Generalized Linear Model in SPSS was used for TEE and RER analysis, which included adjustment for fat-free mass and fat mass. For all analyses, power was set at 80%, with a p value of <0.05.

## Results

### SG is associated with degeneration of CSMG neurons

Confirming our hypothesis, we observed signs of degeneration in CSMG neurons of SG mice 4 weeks post-surgery. This was evidenced by the presence of chromatolysis in CSMG tissue sections stained with NeuroTrace™, a fluorescent Nissl that specifically labels neurons, and imaged with a confocal microscope (figure 1B). No evidence of chromatolysis or other signs of neuronal degeneration were observed in CSMG neurons of SH-AL or SH-IF mice (figure 1B and 1C). During chromatolysis, there is dissolution/loss of the Nissl substance (rough endoplasmic reticulum or RER), nuclear eccentricity, nuclear, nucleolar, and cytoplasmic swelling (figure 1D) (29). A target cell (or cells) innervated by the degenerated neuron loses its innervation unless that neuron regenerates (figure 1E). To further substantiate our hypothesis that SG leads to degeneration of CSMG neurons due to damage to their sympathetic axons innervating the stomach, we performed a neuronal cell count of whole tissue cleared CSMG from SG and sham mice. We used the software Stereology™ as it captures all cells within tridimensional frames, more precisely estimating the total neuronal number. We found that SG mice had significantly fewer CSMG neurons than SH-AL mice, suggesting that after SG, a subset of degenerated neurons undergoes apoptosis (figure 1F). Cell counts were normalized for area and volume (supplemental figure 1). The tissue cleared CSMG were labeled for TH and imaged with a light sheet microscope (figure 1G).

#### CGX recapitulates the weight loss effects of SG in both obese male and female mice

To test whether loss of CSMG neurons would mimic the metabolic benefits of SG, we set out to investigate the metabolic effects of CGX on obese mice. Percent weight loss was similar for male SG and CGX mice, and significantly lower than both the SH-AL and SH-IF groups in the first 3 weeks (Fig.2A) (For a and b, p<0.0001 on week 1, 2, 3 and 4. For a and c, p<0.05). By week 4, SH-IF caught up with the percent weight loss of SG and CGX mice, while SH-AL continued to have a significantly lower percent weight loss. Percent Fat Mass (FM) loss followed a similar pattern as percent weight loss, indicating that weight loss in SG and CGX mice was at the expense of FM loss (figure 2B) (For a and b, p <0.05 for week 1; p<0.001 for week 2; p<0.0001 for week 3, p<0.05 for week 4. For a and c, p<0.001). But by week 4, CGX had a more pronounced FM loss than SG mice. In terms of preservation of lean mass, CGX and SG reversed their initial lean mass (LM) loss, while SH-IF experienced increasing levels of LM loss throughout the study (figure 2C) (For a and b, p<0.0001 for week 2 and 3; and p<0.001 for week 4). SH-AL did not lose LM at any point (figure 2C). Two-way ANOVA with multiple comparisons was used for these analyses and the column factor for all of them was F (3, 127) = 48.20.

**Figure 2.**
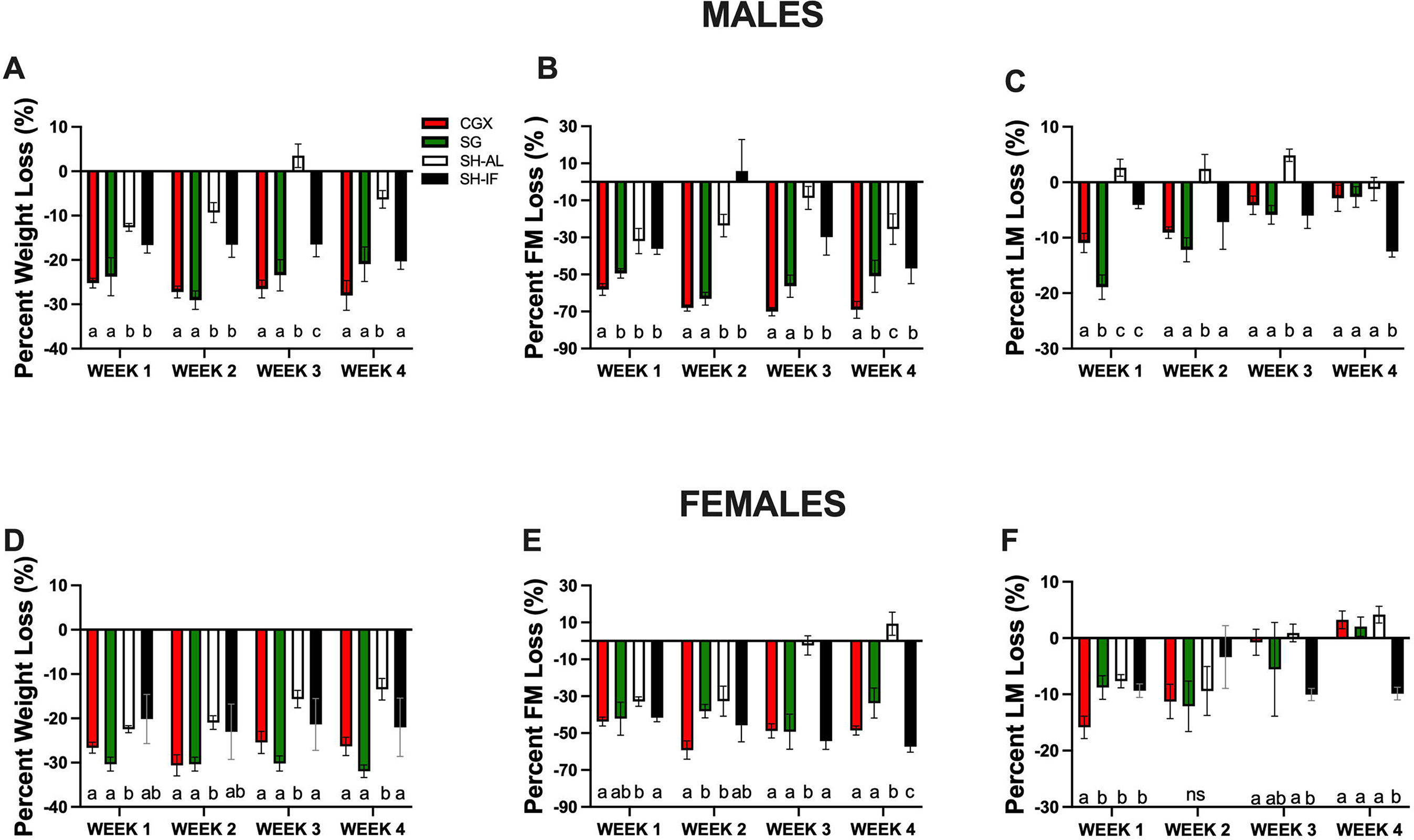
Body weight and body composition in males and females after the four respective interventions: celiac ganglionectomy (CGX), sleeve gastrectomy (SG), sham surgery with ad libitum feeding (SH-AL), and sham surgery with caloric restriction and intermittent fasting (SH-IF). **A**. Male Percent Weight Loss from baseline. For a and b, p<0.0001 on weeks 1, 2, 3 and 4. For a and c, p<0.05. Two-way ANOVA with multiple comparisons, column factor F (3, 127) = 58.75. **B**. Male Percent Fat Mass Loss from baseline. For a and b, p <0.05 for week 1; p<0.001 for week 2; p<0.0001 for week 3, p<0.05 for week 4. For a and c, p<0.001. Two-way ANOVA with multiple comparisons, column factor F (3, 127) = 48.20 **C**. Percent Lean Mass Loss from baseline. For a and b, p<0.05 and a and c, p<0.0001 for week 1. For a and b, p<0.0001 for week 2 and 3; and p<0.001 for week 4. Two-way ANOVA with multiple comparisons, column factor F (3, 127) = 48.20. N=7-14 for each group. **D**. Male weekly energy intake post-surgeries. For a and b, p<0.01 for week 1, p<0.0001 for week 2, and p<0.05 for week 4. For a and c, p<0.01 for week 1, p<0.05 for week 2 and 4. For a and d, p<0.001 for week 2 and p<0.0001 for week 4. Two-way ANOVA with multiple comparisons, column factor F (3, 104) = 31.98. **E**. Female Percent Weight Loss from baseline. SG versus SH-AL, p<0.01; CGX versus SH-AL, p<0.05 for week 1. SG versus SH-AL, p<0.001 and CGX versus SH-AL, p<0.01 for week 2. SG versus SH-AL, p<0.001 and CGX versus SH-AL, p<0.05 for week 3. SG versus SH-AL, p<0.0001 and CGX versus SH-AL, p<0.01 for week 4. Two-way ANOVA with multiple comparisons, column factor F (3, 124) = 7.323. **F**. Female Percent Fat Mass Loss from baseline. CGX versus SH-AL, p <0.01 for week 1; CGX versus SH-AL, p <0.05 for week 2; CGX versus SH-AL, p <0.0001 for weeks 3 and 4. Two-way ANOVA with multiple comparisons, column factor F (3, 128) = 30.46. **G**. Female Percent Lean Mass Loss from baseline. CGX versus SH-AL, p <0.001 for week 1; no significance on week 2; CGX versus SH-IF, p <0.01 for week 3; CGX versus SH-IF, p <0.0001 for week 4. Two-way ANOVA with multiple comparisons, column factor F (3, 128) = 2.217. N=6-14 for each group.

As in males, percent weight loss was significantly greater in female CGX and SG than in SH-AL mice (figure 2D) (a versus b, p< 0.01 for week 1, p<0.001 for week 2, p<0.001, p<0.05 for week 3, and p<0.0001 for week 4; Two-way ANOVA with multiple comparisons, column factor F (3, 124) = 7.323, N=6-14 per group). CGX, SG and SH-IF mice lost significantly more FM than the SH-AL group, especially after the second week post-surgery (figure 2E) (a versus b, p<0.01 for week 1, p<0.05 for week 2, p<0.0001 for weeks 3 and 4; a versus c, p <0.05 for week 4; Two-way ANOVA with multiple comparisons, column factor F (3, 128) = 2.217. N=6-14 per group). While initially all groups lost LM, the SH-IF group was the only one that experienced a profound loss of LM (figure 2F) (a versus b, p<0.001 for week 1; p<0.01 for week 3 and p<0.0001 for week 4; Two-way ANOVA with multiple comparisons, column factor F (3, 128) = 2.217. N=6-14 per group).

In summary, CGX produces similar or greater percent body weight loss and fat mass loss as SG, while preserving lean mass.

#### Weight loss in male CGX and SG mice is associated with high levels of non-esterified fatty acid in the stools

Next, we aimed to find out whether the weight loss experienced by CGX and SG mice was secondary to decreased energy intake, malabsorption, or increased energy expenditure, or a combination of these. Energy intake for male CGX mice was similar to the intake of SH-AL mice at all time points (figure 3A). Male SG mice had low energy intake in the first two weeks post-surgery, as previously described (25). However, after the second post-operative week, male SG mice consumed as many calories as CGX and SH-AL mice. In order to maintain a similar percent body weight and fat mass loss, SH-IF mice had to consume significantly fewer calories than CGX and SG mice (a versus b, p<0.0001 for week 1, p<0.001 for week 2 and 3, and p<0.0001 for week 4; Two-way ANOVA with multiple comparisons, column factor F (3, 133) = 45.21; N=8-13. Energy intake was lower for the female CGX and SG groups compared to SH-AL mice only in the first week post-surgery (figure 3C). Throughout the remainder of the study, all groups except for the SH-IF mice, consumed similar amounts of calories, at times with either CGX or SG consuming more calories than SH-AL mice (figure 3C) (a versus b, p<0.05 for the week 1, p<0.01 for week 2, p<0.05 for week 3, p<0.01 for week 4; a versus c, p<0.05 for week 1, p<0.01 for week 2, p<0.0001 for weeks 3 and 4; Two-way ANOVA with multiple comparisons, column factor F (3, 133) = 45.21; N=8-13).

**Figure 3.**
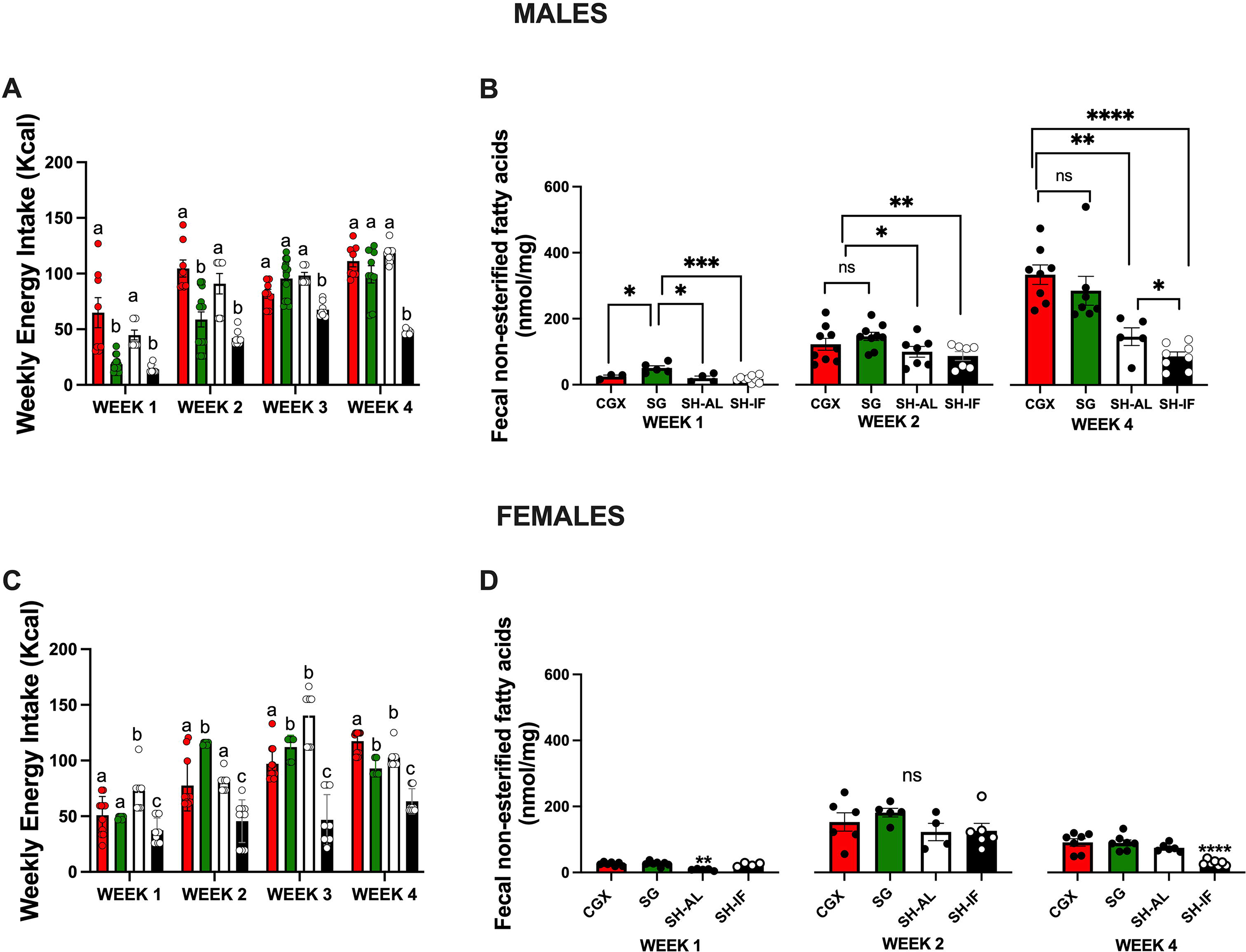
**A**. Male weekly energy intake post-surgeries. CGX versus SG, p<0.0001, CGX versus SH-AL, p>0.05, CGX versus SH-IF, p<0.001 for week 1; CGX versus SG, p<0.001, CGX versus SH-AL, p>0.05, CGX versus SH-IF, p<0.001 for week 2; CGX versus SG, p>0.05, CGX versus SH-AL, p>0.05, CGX versus SH-IF, p<0.01 for week 3; and CGX versus SG, p>0.05, CGX versus SH-AL, p>0.05 and CGX versus SH-IF, p<0.001 for week 4. Two Way ANOVA column factor p<0.0001, F(3,137)=51.75. **B**. Fecal non-esterified fatty acid content at weeks 1, 2 and 4 in males that underwent CGX, SG, SH-AL or SH-IF. **C**. Female weekly energy intake post-surgeries. CGX or SG versus SH-AL or SH-IF, p<0.05 on week 1, with SH-AL consuming more and SH-IF consuming fewer calories than CGX and SG. CGX versus SG or SH-IF, p<0.01, with SG consuming more energy and SH-IF consuming less energy than CGX on week 2. CGX versus SG, p<0.05 and CGX versus SH-IF, p<0.0001 for week 3. CGX versus SG, p<0.001, CGX versus SH-AL, p<0.05, and CGX versus SH-IF, p<0.0001 for week Two-way ANOVA with multiple comparisons, column factor F (3, 133) = 45.21. **D**. Fecal non-esterified fatty acid content at weeks 1, 2 and 4 in females that underwent CGX, SG, SH-AL or SH-IF. One Way ANOVA. N=5-8 mice per group; for ** p<0.01; and **** for p<0.0001.

To find out whether malabsorption could play a role in the weight loss after CGX or SG, we measured the concentration of non-esterified fatty acids in the stool. In males, non-esterified fatty acid stool content was similar across the groups only on week 1, as both SG and CGX mice had significantly higher levels of stool non-esterified fatty acid content in weeks 2 and 4 (figure 3B) (One-way ANOVA. N=7-9; * for p<0.05, ** p<0.01, *** p<0.001 and ****p<0.0001). We did not observe a similar pattern in female mice, except for the SH-IF group having the lowest fecal concentration of non-esterified fatty acids (figure 3D). Female CGX, SG and SH-AL had similar non-esterified fatty acid stool content (One-way ANOVA. N=7-9; ** p<0.01, and ****p<0.0001).

Indirect calorimetry revealed that fat-free mass (lean mass) adjusted total energy expenditure was similar across all groups in males and females (figure 4A and D, nonsignificant, generalized linear model, SPSS). In the male group, SH-AL mice had the lowest RER value (p=0.02, generalized linear model, SPSS), while in the female cohort, CGX mice had the lowest RER (p=0.004, generalized linear model, SPSS). RER values closer to 0.75 indicate that lipid oxidation is favored as opposed to carbohydrate utilization. Locomotor activity was also similar across the four groups both in the male and female cohorts (figure 4C and F) (nonsignificant, Two Way ANOVA with generalized linear model).

**Figure 4.**
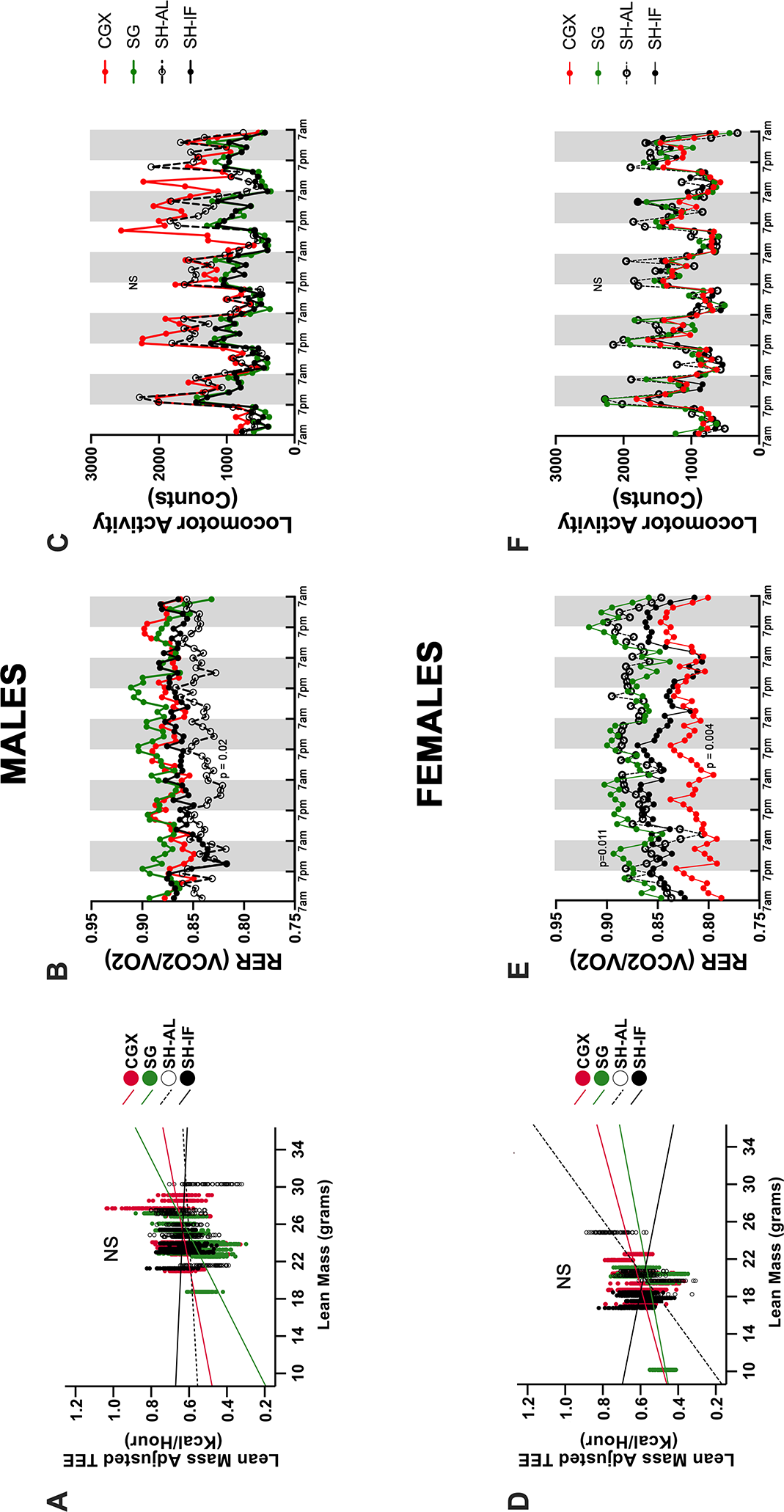
**A and D**. Lean Mass-Adjusted Total Energy Expenditure (TEE) in males and females, respectively. Non-significant differences among CGX, SG, SH-AL and SH-IF groups. Generalized Linear Model in SPSS. **B and E**. Respiratory Exchange Ratio (RER) in males and females, respectively. For males, SH-AL compared to the other groups had the lowest RER, p=0.02. Generalized Linear Model in SPSS. For females, CGX had the lowest RER compared to the other groups, p=0.004, while SG had the highest RER. Generalized Linear Model in SPSS**. C and F**. Locomotor activity (total counts including axes X, Y and Z of the chambers) in males and females, respectively. Non-significant for both male and female cohorts. Two-Way ANOVA with linear modeling.

In summary, a decrease in food intake or an increase in energy expenditure are clearly not the driving forces behind the weight loss after CGX or SG. In males, an increased excretion of lipids in stools may be a contributing factor to the weight loss, while in females that does not appear to be as important. And as previously described for SG mice(25), sham mice required stringent calorie restriction to achieve the same percent body weight loss and fat mass loss as CGX or SG mice.

#### SG and CGX lead to a similar improvement in glucose homeostasis in males

Baseline OGTT was similar for all groups in the female and male cohorts (supplemental figure 1, Two-way ANOVA with multiple comparisons with Tukey test, for baseline p>0.05). Glucose tolerance improved remarkably in the first week post-surgery for male SG, CGX and SH-IF mice compared to SH-AL mice and continued to be better at week 3 (figure 3A and C, One-way ANOVA was used for analysis of AUC. For a versus b, p<0.001 for week 1, and p<0.05 for week 3; a versus c, p<0.05 for week 3). Insulin sensitivity for male CGX and SG mice was better than SH-AL on week 2 but the difference disappeared by week 4 (One-way ANOVA was used for analysis of AUC. For a versus b, p<0.01 and a versus c, p<0.01 for week 2; a versus b, p<0.001 for week 4). Male SH-IF mice had the best glucose tolerance and insulin sensitivity overall. To further characterize the improvement in glucose homeostasis after CGX and SG, we performed a mixed meal test to measure the incretin response 2 weeks post-surgery (figure 5E and 5F). CGX mice had a significantly elevated baseline (fasting) level of serum active GLP-1 compared to SG and controls (One-way ANOVA, p<0.05 for CGX versus SG, and p<0.01 for CGX versus SH-AL and SH-IF). Serum insulin was significantly higher in SH-AL mice, at baseline, in the fasting state. At the 30-minute time point, active GLP-1 continued to be more elevated in CGX and SG than in the controls (p<0.05), while there was no difference in serum insulin levels. A glucose-stimulated insulin secretion (GSIS) test 2 weeks post-surgery revealed that while there is a significant surge in serum insulin concentration after an oral glucose gavage in SG mice, the other mice, including the CGX group, experienced a milder increase in serum insulin (figure 5G) (One way ANOVA, **** for p<0.0001). Supplemental figure 3 shows the blood glucose levels during the GSI, with AUC and fasting blood glucose levels at baseline.

**Figure 5.**
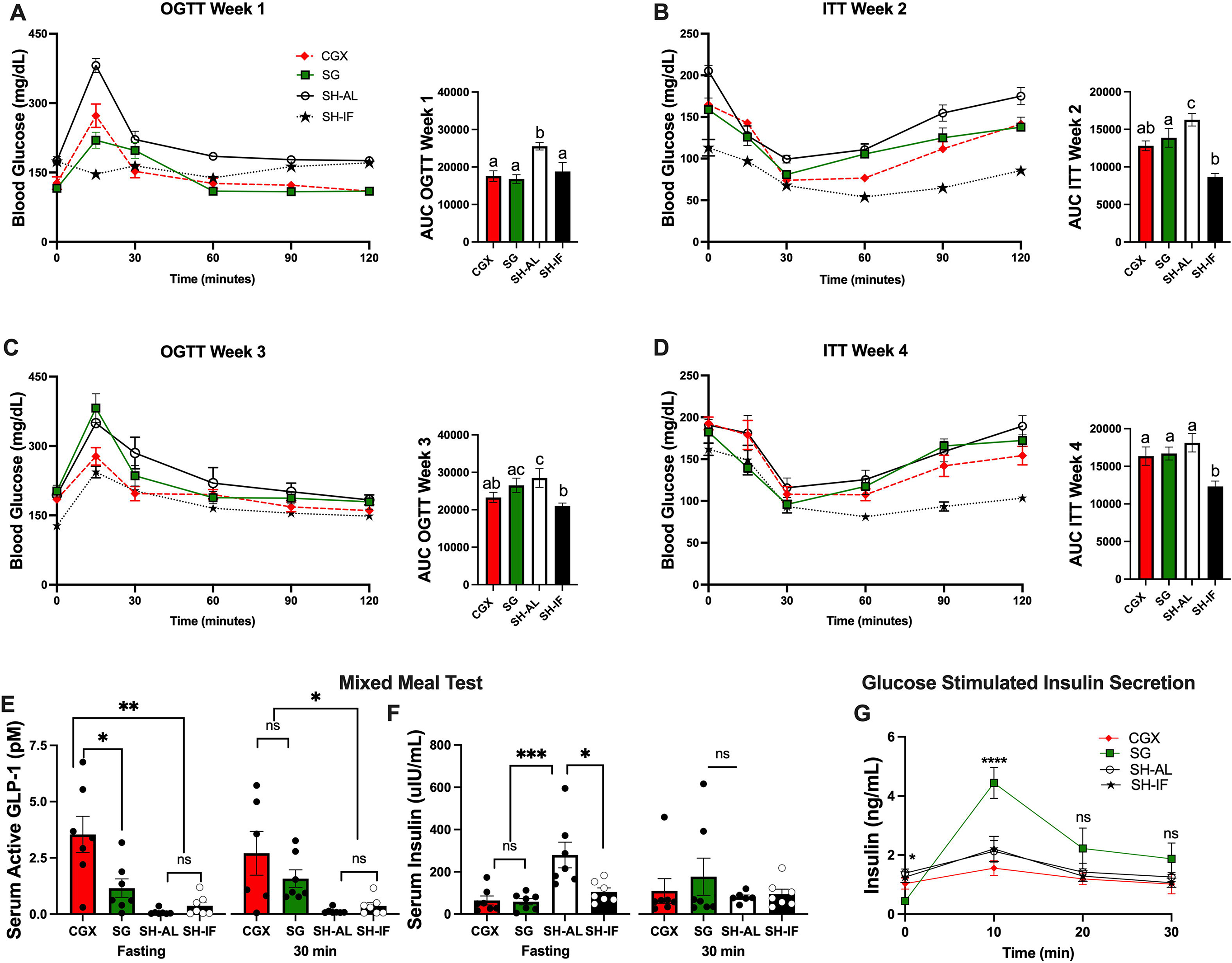
Glucose homeostasis in males. **A and C**. Oral glucose tolerance test on weeks 1 and 3 post-SG, CGX and sham surgery with ad libitum feeding (SH-AL) or with intermittent fasting with caloric restriction (SH-IF). **B and D**. Insulin tolerance test on weeks 2 and 4. One-way ANOVA was used for analysis of AUC. For **A**, a versus b, p<0.001; for **B**, a versus b or c p<0.0001; for **C**, a versus b or c, p<0.05; for **D**, a versus b, p<0.01. N was 8-14 per group, with experiments repeated using different cohorts of mice. **E and F**. Mixed Meal test with oral gavage of Ensure with pre and post (30-minute) values of serum active GLP-1 and insulin. One-way ANOVA. N=7 mice per group. * for p<0.05; ** for p<0.01. **G**. Glucose-stimulated insulin secretion. Insulin levels at time-points zero, 10, 20 and 30 minutes. * for p<0.05; **** for p<0.0001.

**Figure 6.**
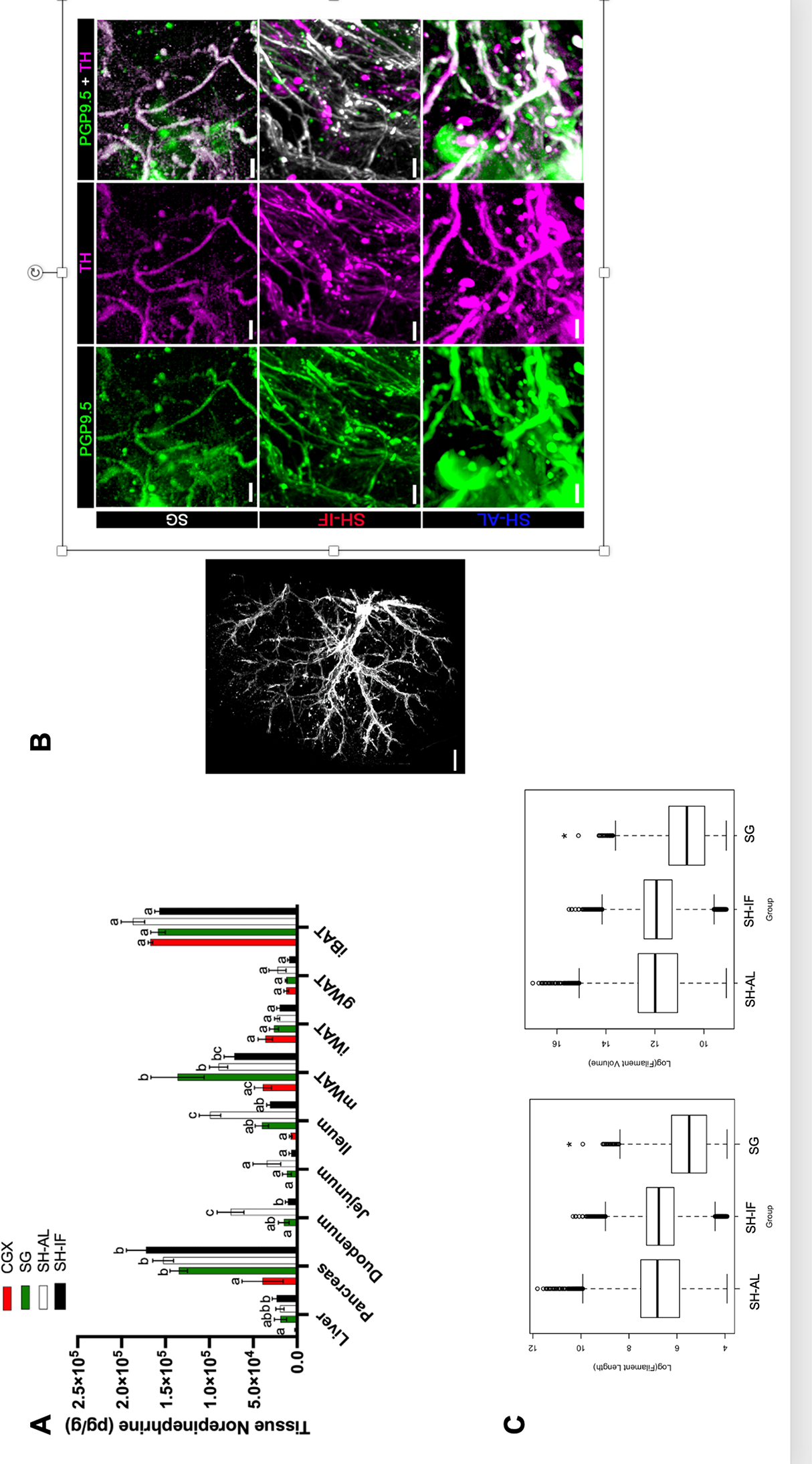
Liver sympathetic innervation. **A**. Tissue norepinephrine content by high performance liquid chromatography at 4 weeks post CGX, SG and sham surgery. Two-way ANOVA with multiple comparisons. N=5 per group. For liver, a and b, p<0.05; for pancreas, a and b, p<0.01; for duodenum, a and b, p<0.05, and a and c, p<0.001; for jejunum, p>0.05; for ileum, a and b, p<0.01, and for a and c, p<0.001; for mWAT, a and b, p<0.01, and a and c, p<0.05; for iWAT, gWAT and iBAT, p>0.05. mWAT – mesenteric white adipose tissue; iWAT-inguinal white adipose tissue, gWAT-gonadal white adipose tissue, iBAT-interscapular brown adipose tissue. **B**. Liver iDisco. In black-and-white, whole liver, scale bar 500 μm. Panel shows PGP9.5 (green), TH-labeled neurons (magenta), and the two markers merged (white). Scale bar 10 μm. **C**. Mixed effects model analysis of filament volume and length for sympathetic axons in liver of mice 4 weeks after SG, SH-AL and SH-IF. p<0.05.

Given that CGX mice experienced a robust improvement in glucose homeostasis that did not appear to be insulin dependent, we measured urine glucose with a chemical dipstick in the light of the fact that the CSMG also innervates the kidneys. Our goal was to assess whether there was an increased rate of urinary glucose excretion in CGX mice. However, urine dipstick testing did not reveal any differences in urinary glucose among the 4 different groups in the male cohort (supplemental Table 1).

Female CGX, SG and SH-IF mice, on the other hand, only experienced improvement in glucose tolerance in the first week post-surgery (supplemental figure 4A, One Way ANOVA, p<0.05). No difference in glucose tolerance or insulin sensitivity was observed in weeks 3, and 4, respectively (supplemental figure 4B and C).

In summary, the improvement in glucose homeostasis after CGX, SG or SH-IF in young adult mice was sex-dependent, as males experienced a more robust improvement than females, unlike what has been described in middle-aged mice(25).

#### CGX and SG are associated with liver sympathetic denervation

As we hypothesized that sympathetic neuronal degeneration in the CSMG partially mediates the weight loss and glycemic effects of SG, we measured tissue norepinephrine content as a surrogate for sympathetic denervation (figure 5A)(Two-way ANOVA with multiple comparisons was used for the analysis. N=5 mice per group. For liver, a and b, p<0.05; for pancreas, a and b, p<0.01; for duodenum, a and b, p<0.05, and a and c, p<0.001; for jejunum, p>0.05; for ileum, a and b, p<0.01, and for a and c, p<0.001; for mWAT, a and b, p<0.01, and a and c, p<0.05; for iWAT, gWAT and iBAT, p>0.05). We measured norepinephrine content in liver, pancreas, duodenum, jejunum, ileum, mWAT, iWAT, gWAT and iBAT. We included mWAT because our tracing studies uncovered that CSMG neurons innervate that adipose tissue depot (supplemental figure 5). Although the CSMG do not innervate iWAT, gWAT, or iBAT, we included those tissues as their sympathetic innervation could have been affected through indirect ways. The data from the tissue norepinephrine content suggested that in male mice, sympathetic denervation of the liver, duodenum, and ileum were common features shared by CGX and SG mice. SG’s liver norepinephrine content was in-between the level for CGX and the level for control mice. SH-IF mice, who towards the end of the period of study observation had lost a significant amount of weight, overall, also displayed decreased tissue norepinephrine content compared to SH-AL mice, who did not lose any significant amount of weight. Regarding mWAT, the CGX group had the lowest tissue norepinephrine content. No significant findings were associated with iWAT, gWAT or iBAT in any groups.

Because of the importance of the liver in maintaining systemic glucose homeostasis, we further sought to determine whether SG was associated with a decrease in hepatic sympathetic axonal density. Using iDisco tissue clearing we found that SG mice had lower filament length and filament volume, compared to SH-AL and SH-IF mice (figure 7B and C) (Mixed model and mixed effects model analysis, p<0.05 for both length and volume, normalized by area). In figure 7B, SG images show a low density of axons, compared to SH-AL and SH-IF mice, observed in PGP9.5 labeling, TH-labeling, and the merging of both.

## Discussion

This is the first study to demonstrate the effects of SG on the CSMG, the major prevertebral sympathetic ganglia innervating the gastric target of this bariatric procedure. Chromatolysis and a decrease in neuronal number in the CSMG of SG mice was not simply the result of laparotomy or abdominal manipulation, as mice that underwent sham surgery had no evidence of neuronal degeneration in their CSMG. We are not the first group to demonstrate that resection of stomach tissue can lead to neuronal degeneration of the sympathetic ganglia supplying its innervation. Gerke (30) demonstrated that in dogs that undergo total gastrectomy, chromatolysis and apoptosis of CSMG neurons can be shown up to one year following the surgery. Ongoing studies aim to demonstrate the presence of apoptotic neurons in the CSMG from SG mice.

The main goal of the present study was to find out whether CSMG neuronal degeneration in the context of SG has biological and clinical relevance. Based on the finding that CSMG ablation (CGX) in obese mice recapitulates the glycemic and weight loss effects of SG in male mice suggests that neuronal degeneration is important for SG’s metabolic effects. A recent study that investigated the role of cannabinoid signaling in the weight loss and glycemic effects of RYGB found that splanchnic denervation causes weight loss and blood glucose improvement in obese mice (31). It is not known how splanchnic denervation and CGX differ, since the splanchnic nerve carries CNS input to CSMG neurons. It is likely that CGX is more effective in causing widespread sympathetic denervation in the abdominal organs it innervates (32). Importantly, the present results support the idea that inhibition of sympathetic drive in obesity appears to alleviate hyperglycemia and promote weight loss.

CGX has been shown to decrease arterial pressure in rodent models of hypertension, underscoring the role CSMG neurons play in regulating cardiovascular function, as they supply innervation to the splanchnic circulation (33, 34). This finding further suggests involvement of CSMG neurons in states of SNS overactivity, such as hypertension. We now demonstrate that ablation of CSMG neurons also ameliorates another state of SNS overactivity, which is obesity with hyperglycemia. CGX in obese mice led to weight loss and fat mass loss that was equivalent to those achieved through SG in both males and females. This loss did not appear to be secondary to any significant effects of CGX on energy intake or energy expenditure. However, it is important to point out that CGX did not lead to metabolic adaptation, as defined by a loss of fat free mass that is the harbinger of a reduction in energy expenditure and weight regain. Metabolic adaptation is a hallmark of diet-induced weight loss in humans and rodents(35, 36), and is partly attributable to a decline in skeletal muscle energy expenditure, occurring even after modest weight loss, defined as less than a 10% loss of body weight (37, 38).

Results from the present study suggest that weight loss in CGX and SG is importantly related to increased fecal calorie excretion in males, as evidenced by the high free fatty acid content in the stools of CGX and SG mice. This points to a malabsorptive phenotype in both CGX and SG mice, consistent with the changes in sympathetic innervation in the duodenum and ileum that we found in the present study. Future studies need to address whether the fat malabsorption phenotype we found is secondary to bile acid malabsorption, impaired nutrient absorption, or microbiome changes. Although SG is not classically viewed as a malabsorptive procedure, recent studies have revealed that SG in humans is associated with evidence of malabsorption of oral anticoagulants and birth control pills (39, 40).

CGX in obese mice was also associated with improved glucose homeostasis in males. It is not clear whether CGX’s effects on blood glucose are weight-loss independent. CGX led to increased serum levels of active GLP-1, although, unlike SG, the levels were already elevated at baseline, when the mice were fasting. Future studies will need to evaluate whether increased GLP-1 secretion is a mechanism through which CGX influences glucose homeostasis, independently of CGX-induced weight loss. Other potential mechanisms for CGX to ameliorate blood glucose independent of weight loss include liver, pancreas and mWAT sympathetic denervation, which the low norepinephrine content in those tissues suggest may be important contributing factors. Neurons in the CSMG are the main source of adrenergic efferent projections to the liver (9, 10). Autonomic input descending from the hypothalamus and transmitted through efferent projections in the periphery regulates liver glycogen metabolism(41). Increased sympathetic outflow to liver results in an increase in hepatic glucose output(42). This occurs due to the stimulation of hepatic glycogen phosphorylase and phosphoenolpyruvate carboxykinase, which promote glycogenolysis and gluconeogenesis, respectively(43). The pancreas also receives adrenergic innervation that originates in the CSMG(8). When the CSMG is resected in rats, only 10% of adrenergic axons are left in the pancreas, confirming that the CSMG is the main source of pancreatic sympathetic innervation(11). The pancreatic islets are abundantly innervated, as first recognized by Langerhans(44). While the pancreatic parasympathetic innervation in humans, compared to mice, is sparser, adrenergic fibers richly innervate arterioles, capillaries, and the islets in both humans and rodents(45). Stimulation of α adrenergic receptors on pancreatic beta cells leads to potent inhibition of insulin secretion(46, 47). On the other hand, beta adrenergic receptor stimulation on pancreatic alpha cells promotes glucagon secretion(48). Despite significant advances in the understanding of hepatic and pancreatic sympathetic innervation, little is known about whether there is a direct role for CSMG neurons in metabolic regulation or whether specific CSMG neuronal populations are specialized in the regulation of hepatic or pancreatic function. Regarding mWAT function, it is the only visceral adipose tissue depot that is equivalent in humans and rodents(49). Using Pseudorabies virus tracing, Nguyen *et al* found that mWAT is innervated by the sympathetic chain ganglia (T5-L3)(50). However, in that study, the CSMG was not checked for labeling. In the present manuscript, we report that we found that CSMG neurons innervate mWAT. Moreover, we found that CSMG neurons projecting to liver, pancreas and mWAT are mostly specific, rarely co-localizing.

What further suggests that the glycemic benefits conferred by CGX in males are weight-independent is the fact that although female CGX mice experience a similar rate of weight and fat mass loss, the same cannot be said about their glucose homeostasis. Our experience with middle aged female DIO mice revealed that they achieve blood glucose lowering after SG or SH-IF(25). Since the female mice in the present study were significantly younger than the 45–48-week-old mice that we previously studied, we wonder if their higher glucose intolerance and insulin resistance was age- and strain-related. We tested two consecutive cohorts of female DIO mice at 18 weeks of age arriving at the same results. Further testing with different strains of obese female mice will help us understand this puzzling finding. Additionally, female CGX and SG mice did not have the same degree of lipid stool excretion as male CGX and SG mice, suggesting that the underlying weight loss mechanisms after CGX or SG may be sexually dimorphic.

Our study findings raise many questions regarding what is shared by CGX and SG in terms of mechanisms of weight loss and blood glucose lowering. Future studies need to focus on how CGX promotes metabolic benefits in obese mice and the nature of the sex differences found in some of its effects. CSMG neuronal degeneration as a potential mediator of some of SG’s metabolic benefits is an exciting finding as it could pave the way for the development of new therapies to treat obesity and type 2 diabetes, as recently published(51).

## Funding

Berrie Neuroscience of Obesity Initiative at Columbia University - ABE

7K08DK101830-05 – ABE

Berrie Foundation Fellowship Award – MK

## Contribution Statement

GJS and ABE were responsible for the project conceptualization, design, data acquisition, statistical analysis, and manuscript preparation. ROB, NRL, MK, CSJ and AD were involved in data acquisition and interpretation.

## Figure Legends

**Supplemental figure 1**. **A**. Total neuronal cell count in the celiac superior mesenteric ganglia (CSMG) of sleeve gastrectomy (SG) and sham surgery with ad libitum feeding (SH-AL) mice. Unpaired student’s t test, p<0.05. **B**. Total CSMG area (μm^2^) in SG and SH-AL mice. Unpaired student’s t test, non-significant. **C**. Total CSMG volume (μm^3^) in SG and SH-AL mice. Unpaired student’s t test, non-significant.

**Supplemental figure 2**. **Baseline OGTT in females and males prior to surgeries**. Two Way ANOVA with multiple comparison, non-significant.

**Supplemental figure 3**. Blood glucose during OGTT, AUC and baseline blood glucose during glucose stimulated insulin secretion test. * for p<0.05; ** p<0.01; and **** for p<0.0001. One Way ANOVA. N=7-11 mice per group.

**Supplemental figure 4**. **Glucose homeostasis in females only**. Oral glucose tolerance test (OGTT) and insulin tolerance test (ITT). Glucose tolerance was improved in CGX, SG and SH-IF compared to SH-AL in week one (p<0.05). No statistically significant difference among CGX, SG, SH-AL and SH-IF for OGTT or ITT on weeks 3 and 4. Two-way ANOVA with multiple comparisons.

**Supplemental figure 5**. **Light sheet microscope images of cleared CSMG labeled with CTb**. **A**. CTb staining of CG neurons projecting to liver (magenta) and pancreas (red). **B**. CTb staining of CG neurons projecting to mesenteric white adipose tissue. **C and D**. CTb staining of CG neurons projecting to liver (blue), pancreas (green) and mesenteric white adipose tissue (pink). Mice were injected in the liver parenchyma of right and left lobes; head, body, and tail of pancreas; and mesenteric white adipose tissue. Note the widespread presence of sympathetic puncta in C and D. Scale bar for all images is 20 μm. N=3 for all studies, obese male mice.

**Table 1**. Detection of urine glucose in males with chemical dipsticks.

