## Supplementary figures and images for "DISRUPTION OF SYMPATHETIC OUTFLOW TO INTRA-ABDOMINAL ORGANS EMULATES THE METABOLIC EFFECTS OF SLEEVE GASTRECTOMY IN OBESE MICE"

### Supplemental figure 1

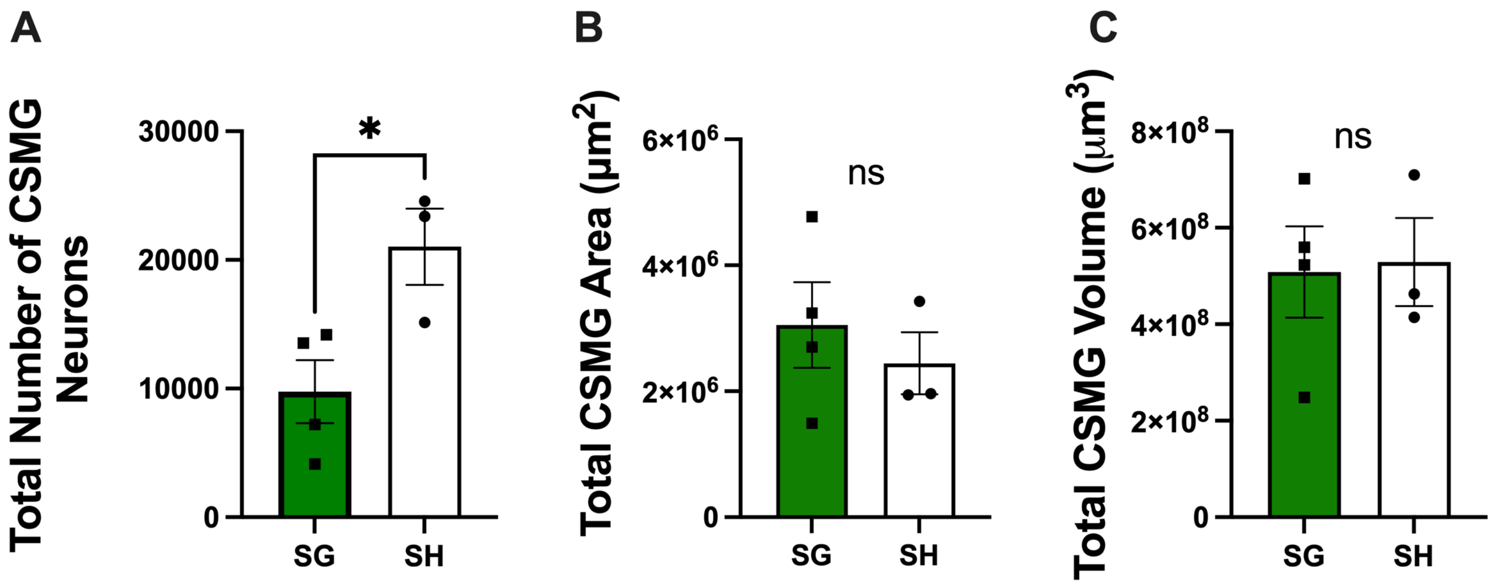

### Supplemental figure 2

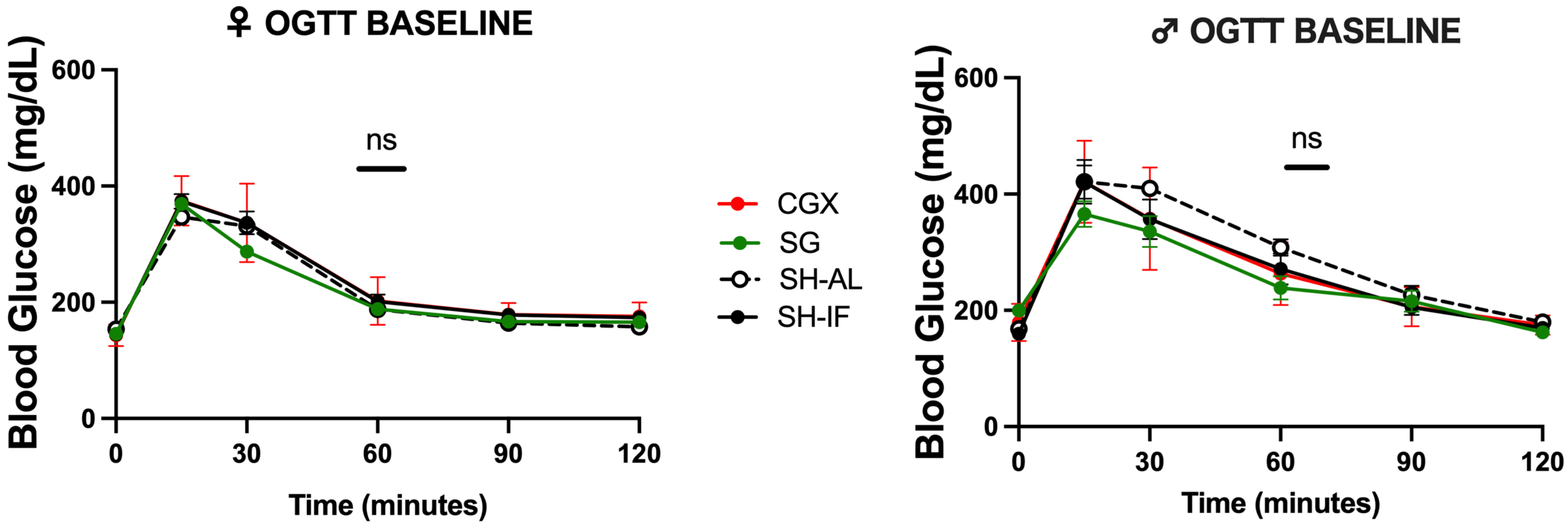

### Supplemental figure 3

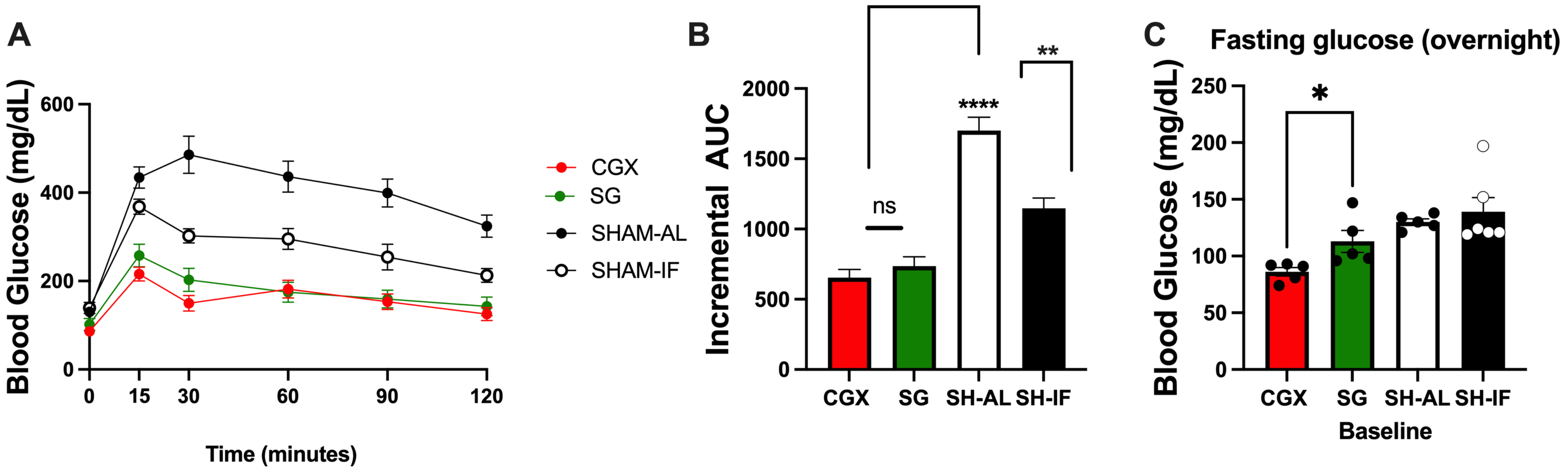

### Supplemental figure 4

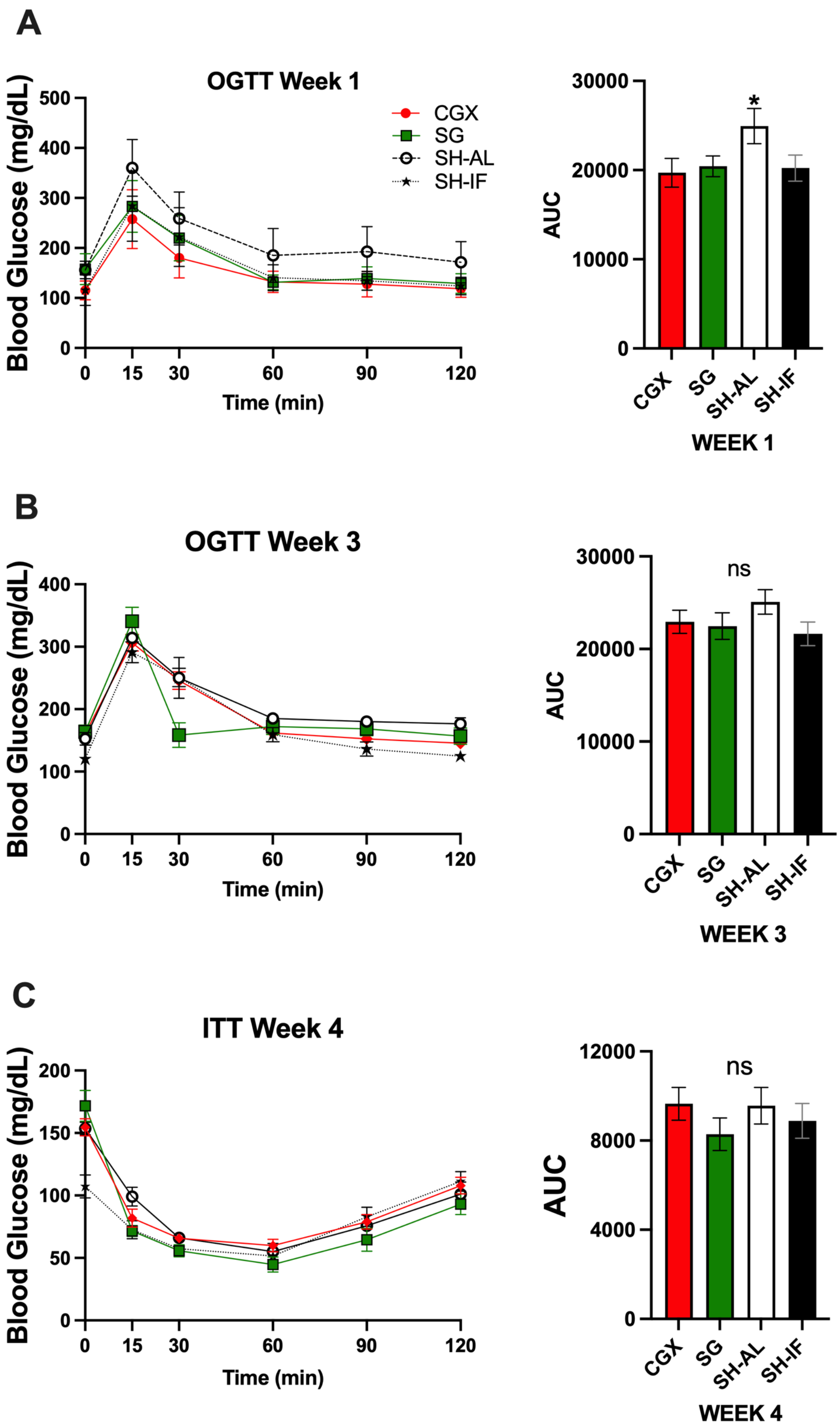

### Supplemental figure 5

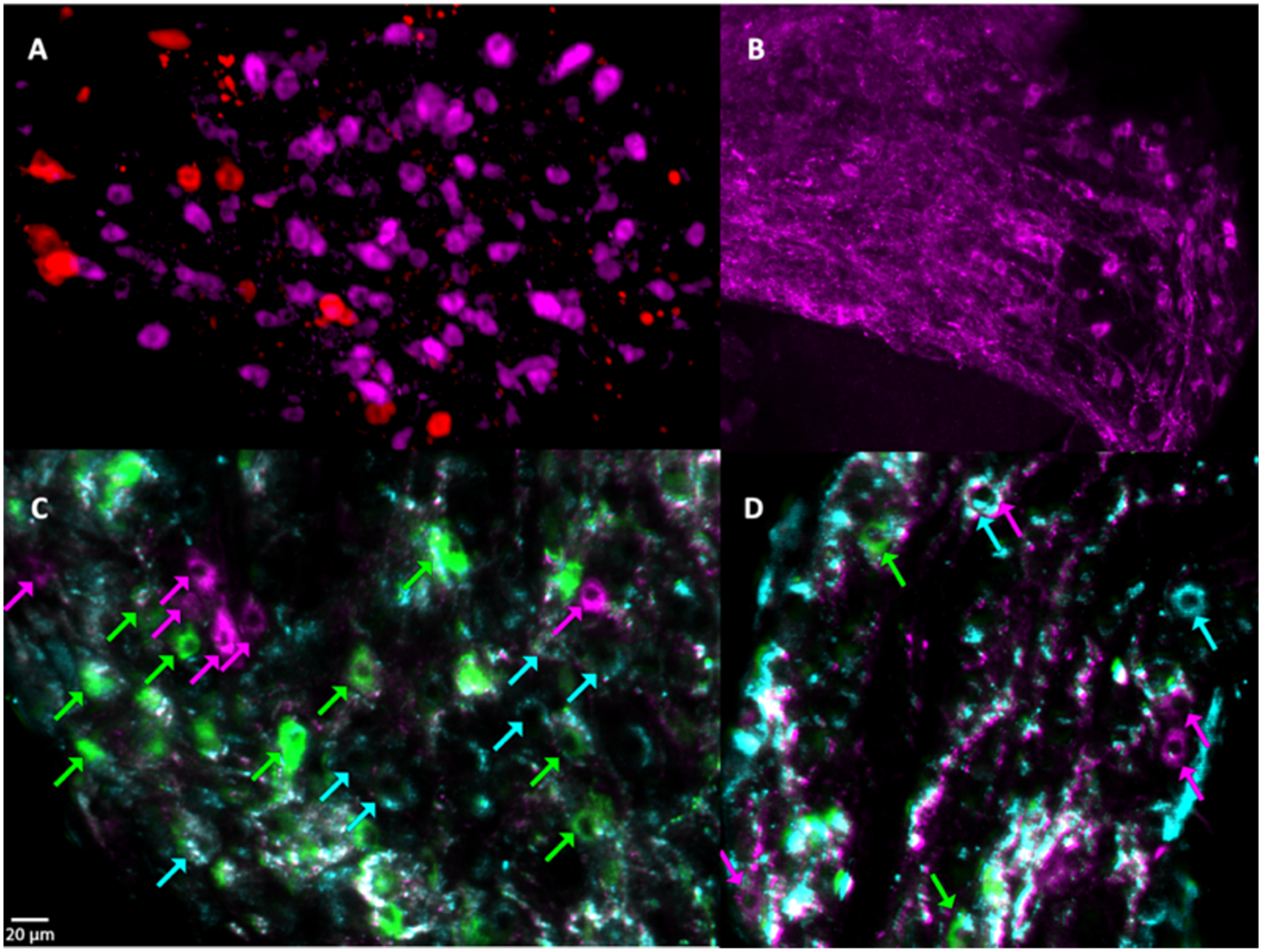
